# Wheat leaf dark respiration acclimates more strongly at night than in the day when responding to nocturnal warming

**DOI:** 10.1101/2025.10.03.680246

**Authors:** Pratima Rana Shahi, Andrew P. Scafaro, Rebecca J. Thistlethwaite, Owen K. Atkin, Richard M. Trethowan, Romina Rader, Adrienne Burns, Onoriode Coast

## Abstract

Rising night temperatures pose a significant threat to wheat productivity, yet the physiological basis of wheat adaptation to nocturnal warming remains poorly understood. We evaluated leaf photosynthetic and respiratory traits in ten Australian wheat cultivars released between 1901 and 2012 to warm nights under temperature-controlled environments. When exposed to warmer nights, rates of leaf net CO_2_ assimilation measured at 25 °C (*A*_net_^25^) remained stable across cultivar release date despite declines in photosynthetic capacity (*V*_cmax_ and *J*_1500_) in newer cultivars. In most cultivars leaf respiratory CO_2_ release in the dark (*R*_dark_) exhibited divergent thermal responses: warm nights suppressed temperature-normalised night *R*_dark_ (*R*_night_) but stimulated or maintained *R*_dark_ in the daytime (*R*_day_). The results suggest that century-long, yield-focused selection may have inadvertently maintained *A*_net_^25^ under warmer nights in modern cultivars through selection for more night-temperature sensitive but efficient photosynthetic capacity (i.e. greater return per protein investment) and overall reduced respiratory demand for maintenance of processes such as Rubisco protein turnover and synthesis. Our findings highlight trait-based targets for enhancing energy efficiency and climate resilience in wheat and opportunities to improve the parameterization of *R*_dark_ to warm nights in crop and Earth system models.

## 1.0 Introduction

Wheat is a staple food crop for about 3 billion people – providing 20% of human calorie and protein intake (Shewry & Hey, 2015). It is cultivated in over 220 million ha worldwide (Hay et al., 2022; Ramirez-Villegas et al., 2020) and demand for wheat is expected to rise as global population increases to 9 billion by 2050 (Alexandratos & Bruinsma, 2012; Kinnunen et al., 2020). Meeting growing demands for wheat is challenging, and is compounded by human-induced climate change (Mondal, 2010; Poveda et al., 2018). Climate change has been marked by increased land surface temperatures at global, regional and national scales compared to pre-industrial levels. The increase in global day and night temperatures have been asymmetric with night time minimum temperature increasing up to 1.4-fold faster than day time maximum temperatures at the global scale (Davy et al., 2017; Jones et al., 1999; Sillmann et al., 2013; Vose et al., 2005; Zhuang & Zhang, 2020). The rise in night temperatures has been attributed to cloud cover, anthropogenic greenhouse gases induced radiative forcing, and vegetation feedbacks (or greening) which reduce radiative heat loss at night (Kaiser, 1998; Shi et al., 2021). Cloud cover further reduces the amount of incoming solar shortwave radiation during the day which lessens daily maximum warming, while also trapping heat leaving the Earth’s surface at night (Karl et al., 1984; Kukla & Karl, 1993). Other minor contributors are land use changes, urban warming effects and soil moisture (Lough, 1995; Shi et al., 2021).

It is predicted that nocturnal warming will continue (Cox et al., 2020; Doan et al., 2022; Wang et al., 2020; Zhuang & Zhang, 2020) if anthropogenic greenhouse gas emissions are not mitigated. Despite night temperatures rising more rapidly than day temperatures, and the well-documented negative effects of warm nights on crop yield (as described below), research in crop thermal physiology has become increasingly asymmetrical with more studies focusing on responses to high day temperatures than to high night-time temperatures (Reynolds et al., 2021). Hot nights pose serious risks to ecosystem functions and services, and crop yields (Sadok & Jagadish, 2020; Welch et al., 2010). Night warming has been strongly linked with wheat yield decline, more so than rising day temperatures (Fischer et al., 2022; García et al., 2015). Wheat grain yield loss of 2–9% °C^−1^ has been reported from field and modelling studies in the USA and Mexico (Fischer et al., 2022; McAusland et al., 2023; Russell & Van Sanford, 2020). A similar response has been reported for rice, with rice grain yield declining 10% for every 1 °C increase in growing-season minimum temperature (Peng et al., 2004). Cereal yield losses due to warm nights are likely due to disrupted reproductive processes (Coast et al., 2014; Narayanan et al., 2018); shortening of the critical development period for determination of grain yield (Fischer et al., 2024), increased respiratory CO_2_ losses during the night (Garcia et al., 2016; Impa et al., 2019) and reduced capacity for carbon fixation through photosynthesis (Coast et al., 2024; Posch et al., 2019). Wheat leaf net CO_2_ assimilation (*A*_net_) and respiratory CO_2_ release (*R*_dark_), which contribute strongly to total carbon budget and ultimately yield, are sensitive to temperature. In the short-term, *A*_net_ increases with instantaneous temperature rise from sub-optimal levels reaching a peak at an optimum temperature before declining with further warming (Posch et al., 2019). This curvilinear response reflects the impacts on photosynthetic CO_2_ fixation, including by reduced Rubisco fixation and electron transport capacity with rising temperature and CO_2_ release by photorespiration and *R*_*dark*_ (Yamori et al., 2014; Scafaro et al., 2024). Over long time frames, warmer growth conditions may cause increases in the thermal optimum of *A*_net_ (Berry & Bjorkman, 1980; Way & Yamori, 2014). However, heat stress including warm nights can cause down-regulation of both *A*_net_ and photosynthetic capacity (i.e. maximum carboxylation rate, [*V*_cmax_] and maximum electron transport rate, [*J*_max_]). For example, inhibition of photosynthesis under warm nights occurs in some wheat cultivars (Posch et al., 2019; Coast et al., 2024), and other crops such as maize (Sinsawat et al., 2004) cotton (Reddy et al., 1997), and soybean (Sankarapillai et al., 2025).

Like photosynthesis, *R*_dark_ responds to instantaneous and long-term temperature change, but the response patterns vary depending on the timeframe being considered. In the short-term, a rise in temperature induces a near-exponential increase in *R*_dark_ (Atkin & Tjoelker, 2003) up to a maximum at around 50-60 °C (T_max_) beyond which respiration rate abruptly declines (O’Sullivan et al. 2017). The quick rise in *R*_dark_ is linked to increased energy demand for upkeeping functions, and diversion of assimilated carbon away from growth (Scafaro et al. 2021). Beyond T_max_, the rapid decrease in *R*_dark_ likely reflects irreversible injury to the respiratory apparatus due to loss of mitochondrial function and a rapid onset of tissue death (O’Sullivan et al., 2013). In the long-term (e.g. days to weeks), higher growth temperature causes a change in energy demand and membrane fluidity which results in reduced temperature-normalised *R*_dark_, reflecting *R*_dark_ acclimation to higher growth temperature. However, limited studies in wheat suggest that CO_2_-based temperature-normalised *R*_dark_ measured during the day (*R*_day_) does not acclimate to night warming. Posch et al. (2022) reported no evidence of acclimation of *R*_day_ measured at 25 °C (*R*_day_^25^) in wheat to warm nights – plants grown under warmer (20 and 25 °C) nights relative to cool (15 °C) nights showed similar *R*_day_^25^. By contrast, CO_2_-based *R*_dark_ during the night (*R*_night_) measured at 25 °C (*R*_night_^25^) acclimates to higher night temperature (Impa et al., 2019). However, these studies were based on a narrow genotypic range – two genotypes in Posch et al. (2022) and six in Impa et al. (2019) – and it remains unclear whether this pattern is consistent across broader wheat germplasm. This highlights the need for further work assessing leaf respiratory responses to night warming across a broader range of wheat cultivars.

Breeding advances and agronomic management have played major roles in wheat yield improvements (Richards et al., 2014; Robertson et al., 2016). Wheat genetic improvement has paralleled enhancement of leaf photosynthesis, for example in China (Tian et al., 2011; Zheng et al., 2011), Mexico (Fischer et al., 2022) and Israel (Blum, 1990). However, it is unclear if enhanced leaf photosynthesis (*A*_net_ measured at a common temperature, e.g. at 25 °C; *A* _net_^25^) or temperature-normalised lower *R*_dark_ would hold under the progressively warmer climate (and warming nights) that emerged over the last century. Considering the drag on wheat yield by high night temperature stress globally, a key challenge to increasing future wheat yield will require deeper understanding of photosynthetic and respiratory adjustments and their links with other leaf functional traits – leaf nitrogen (N) content (Crous et al., 2022; Dusenge et al., 2020; Dusenge et al., 2021; Wang et al., 2020), and leaf mass per unit area (LMA) (Atkin et al., 2006; Manishimwe et al., 2022; Sage & Kubien, 2007) – in historical and modern cultivars to warm nights.

Our study examines 10 historic and current Australian wheat cultivars released between 1901 and 2012 (i.e. over a century of breeding) to understand wheat photosynthetic and respiratory adjustments to warm nights. The cultivars were exposed to cool or warm nights and a standard day temperature in controlled environment facilities to assess wheat leaf photosynthetic traits (i.e. *V*_cmax_, *J*_max_, A_net_), CO_2_-based *R*_dark_ (during the day, *R*_day_, or at night, *R*_night_), and changes in leaf structure and chemical traits (LMA, and leaf N content) under warm nights. We hypothesised that over the last 111 years breeding indirectly selected for the following changes under warm nights: (i) modern cultivars would exhibit higher A_net_^25^ in warm-night-grown plants relative to cool-night controls; (ii) increased leaf photosynthetic capacity (*V*_cmax_ and *J*_max_) underpin higher rates of A_net_^25^; (iii) reduced respiratory CO_2_ losses (lower temperature-normalised rates of *R*_day_ and *R*_night_ in plants exposed to warm nights compared to cool night controls).

## 2.0 Materials and methods

The experiment was conducted in controlled-environment facilities (glasshouses and growth chambers) at the University of New England (UNE) in Armidale, Australia (30.48°S, longitude 151.63°E, elevation 1021.5 m a.s.l.) in 2022.

### 2.1 Plant materials and growth conditions

A set of 10 wheat genotypes (Supplementary Table 1) were used in this experiment. The genotypes included commercial Australian wheat varieties spanning over a century (1901-2012) of breeding. Germplasm for the experiment was sourced from the I. A Watson Grains Centre in Narrabri (NSW Australia 30.27°S, 149.81°E). Seeds of ten Australian wheat cultivars were planted in 4.5 L plastic pots (18 cm diameter and 60 cm height) containing Professional premium potting mix with slow-release fertiliser (J.C. & A.T. Searle Pty. Ltd., Qld. Australia). Potted plants were raised in glasshouse bays set to day/night temperatures of 20/12 °C, 70% relative humidity and a natural photoperiod of 12 hours. The plants were grown for four weeks in the glasshouse until the tillering stage (Zadok 20-29) (Zadoks et al., 1974). During this period, watering was provided manually, and standard operating protocols used for plant husbandry in the glasshouse applied. At tillering, when all plants had a fully extended third leaf, wheat plants were moved into growth chambers (Conviron, Conviron Inc., Winnipeg, Canada), and raised for three weeks till their booting stage (Zadok 30-49).

The three growth chambers were equipped with incandescent and fluorescent lamps, and set to a 14-hour photoperiod with photosynthetic photon flux density (PPFD) of 800-850 µmol m^−2^, and relative humidity of 60%. The experiment was a complete randomised design with two treatments and three replicates. However, each cultivar had six technical replicates (i.e. two pots per chamber per treatment). The treatments were the control temperature (25/12 °C, maximum day/minimum night) and high night temperature (25/22°C, maximum day/minimum night). The experiment was conducted in two sequential batches, with all control plants grown first, followed by all high night temperature plants. Each batch used the same three growth chambers, serving as biological replicates, to ensure consistent environmental conditions across treatments. Plants were watered daily, and pots were kept in the trays containing about 1 cm deep water to avoid water stress. No major pests and diseases were noticed during the experiment. Maximum and minimum temperatures were maintained for 8 hour each in all growth chambers with a transition period of 4 hour between maximum day/minimum night temperatures. Temperature and relative humidity (RH) at canopy level were recorded every five minutes using Tiny-tag Ultra 2 Hasting temperature/RH data loggers (Gemini Data Loggers UK LTD, UK) in all growth chambers during both treatments.

### 2.2 Photosynthesis and respiration

Gas exchange measurements were conducted on recently fully developed flag leaves on tagged main tillers in each treatment using portable infrared gas analyser (IRGA) units (Li-6400XT, LI-COR Inc., Lincoln, NE, USA). The net CO_2_ assimilation (*A*_net_^25^) measurements were taken with Li-6400XT units with 6 cm^2^ leaf chamber and block temperatures set to 25 °C, CO_2_ fixed at 400 ppm with a flow rate of 500 μmol s^−1^ and light intensity of 1500 μmol m^−2^ s^−1^ of Photosynthetically Active Radiation (PAR). Dark-adapted temperature-normalised *R*_day_ and r*R*_night_ were recorded with the same Li-6400XT units. Each chamber’s head block temperature was set to 25 °C for *R*_day_, and 20 °C for *R*_night_. The set chamber temperature resulted in average leaf temperature of 25.5 °C for *A*_net_^25^ and *R*_day_ and 21.3 °C for *R*_night_ (Supplementary Fig. S1). Measurements of *A*_net_^25^ and *R*_day_ (after at least 20 minutes of dark adaptation) were completed during the day (between 09:00 and 17:30 h) and *R*_night_ measured at night (after 20:00 h).

### 2.3 Maximum Rubisco activity and electron transport rate

Two Li-6400XT units were used to assess photosynthetic capacity of the ten wheat genotypes. Photosynthetic CO_2_ response curves (*A:C*_i_ curves) were generated, at constant irradiance of 1500 μmol photons m^−2^ s^−1^, by varying the CO_2_ inside the Li-6400XT leaf chambers as follows: 400, 40, 60, 100, 150, 250, 400, 400, 600, 800, 1000, 1200, 1400 and 400 μmol mol^−1^. Relative humidity within the chamber was maintained between 40 and 70% for all measurements. The maximum Rubisco activity at 25 °C, *V*_cmax_^25^, and electron transport rate at PAR of 1500 μmol m^−2^ s^−1^ and measured at 25 °C, *J*_1500_^25^ were calculated using the leaf biochemical model of photosynthesis (Farquhar et al., 1980) with kinetic constants derived for wheat (Silva-Perez et al., 2018). Response curves of *A*_net_^25^ to the intercellular CO _2_ concentration (*C*_i_) were measured in the mid-section of the flag leaf when the plants reached Zadoks stage (40-49). The relationship between *A* _net_^25^ and CO _2_ followed the FvCB model (Farquhar et al., 1980; von Caemmerer & Farquhar, 1981) with a simple function for limitation by triose phosphate utilization (TPU) (Sharkey et al., 2007). The approach of Gu and Jerome (2010) was used where all possible carboxylation limitation-state combinations were tested, given the required order of limitation states along the C_i_ axis (Rubisco limited <electron transport limited < TPU limited) and the minimum number of data necessary for each limitation state (*n*≥2 when Michaelis Menten constants for Rubisco catalysis of carboxylation, Kc, and oxygenation reactions, Ko; and the photosynthetic CO_2_ compensation point in the absence of mitochondrial respiration in the light, Γ*, are fixed). Representative *A:C*_i_ curves with model fits are presented in Supplementary Fig. S2. The R Language and Environment function optim (R Core Team, 2018) was used to minimize the distribution-wise cost function, and the model with the lowest cost function value was accepted after checking for admissibility and, if necessary, testing for co-limited ‘swinging points’ (Gu & Jerome, 2010). These measurements were taken within a day between 09:00 and 17:30 hr (~40 min before sunset) for each plant.

### 2.4 Determination of leaf mass per area and nitrogen content

Leaf area (cm^2^) was calculated using the same leaf used for gas exchange measurements. Leaves were dried at 65 °C for 72 h or until constant dry weight was achieved then used to calculate leaf dry mass per unit area (LMA, g m^−2^). Then, 1 mg of dry leaf section of each sampled tissue was weighed and analysed to calculate leaf N (%) using an automatic element analyser (Sercon Control “Callisto CF-IRMS” Version 30.0.11, Sercon Integra 2) coupled to a mass spectrometer (Elemental microanalysis LTD, United Kingdom). Using determined N content (%) and LMA, we estimated leaf nitrogen per area (N_area_, g m^−2^) and leaf nitrogen per dry mass (N_mass_, mg g^−1^).

### 2.6 Statistical analysis

All data analyses were conducted in R (R Core Team, 2023). Trait responses to night temperature treatment, cultivar, and their interaction were analysed using analysis of variance (ANOVA) based on a complete randomised design. Data manipulation, visualisation, and statistical modelling were performed using the dplyr, ggplot2, and gridExtra packages. For each trait, means were calculated across cultivars for each treatment and Year of Release.

Chronological trends in physiological traits (e.g., *A* _net_^25^, temperature-normalised *R*_dark_, leaf N content) were assessed using both linear and second-order polynomial regressions. Models were fitted separately to the control and high night temperature datasets, and model selection was based on comparisons of adjusted R^2^ and Akaike Information Criterion (AIC). An F-test was also used to determine whether the polynomial model provided a statistically significant improvement over the linear model. Non-linear models were adopted only when they satisfied all three criteria: higher adjusted R^2^ (by >0.02), lower AIC, and significant F-test (*p* < 0.05).

To enable comparison of *R*_day_ measured at 25 °C (*R*_day_^25^) and *R*_night_ at a common temperature (i.e. at 25 °C), we used the global polynomial respiration-temperature model (GPM) of Heskel et al. (2016) to estimate *R*_night_ at 25 °C (*R*_night_^25^) for each individual leaf *R*_night_ measured at 20 °C (*R*_night_^20^). The GPM model [Eq. 1] is represented below:

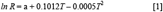

Where *ln R* is natural-log–transformed (*ln*) rate of leaf *R*_night_ at a given leaf *T*, and *a* is a coefficient derived from resolving Eq. 1 given individual leaf *R*_night_^20^ and leaf *T* of 20 °C. The mean estimate of *a* was −2.1443 (95% CI −2.2620, −2.0266) compared to Heskel et al. (2016) with global mean *a* of −2.2276 (95% CI −2.3966, −2.0586). Differences between *R*_day_^25^ and *R*_night_^25^ were tested using two-sample Welch’s t-tests as the measurements were independent observations, samples sizes were unequal, and variances were not assumed to be equal between *R* _day_^25^ and *R*_night_^25^. Unless otherwise stated, results presented for *R*_night_ are of *R* _night_^20^.

To evaluate thermal acclimation across breeding years, derived trait ratios (e.g., *J*_1500_^25^/*V*_cmax_^25^, *R*_day_/*V*_cmax_^25^, and *R*_night_/*V*_cmax_^25^) were calculated and modelled similarly. Where relevant, trait relationships with leaf nitrogen were also explored using regression analysis to assess the coordination between biochemical and physiological traits under control and high night temperature conditions.

## 3.0 Results

### 3.1 Net photosynthesis: no systematic change in over a century of wheat breeding except per unit leaf nitrogen which declined over time

Rates of *A*_net_^25^ per unit leaf area differed among cultivars across both control and high night temperature treatments (*F*_3,114_= 3.05, *p* = 0.031; Figure 1A). While the main effect of Year of Release was not significant when fitted as a continuous variable (*p* = 0.180), a significant interaction between treatment and Year of Release (*p* = 0.006) indicated that cultivar rankings differed between treatments. For instance, the early cultivar Federation (1901) exhibited the highest *A*_net_^25^ (25.5 µmol CO_2_m^2^s^−1^) under control conditions but showed a pronounced decline (19%) under high night temperature (Figure 1a). By contrast, the modern cultivar Suntop (2012) which had modest *A*_net_^25^ under control conditions significantly increased *A*_net_^25^ (24%) under high night temperature (Figure 1a). Overall, there was no significant trend as a function of Year of Release under either treatment (Fig. 1a; both *p* > 0.05) suggesting that breeding over time has not led to systematic change in *A*_net_^25^. The associated stomatal conductance measured at 25 °C of the cultivars and their responses to night temperature were similar to those recorded for *A*_net_^25^ per unit leaf area (Figure 1b). The slopes of the *A*_net_^25^ vs Year of Release plots did not differ between the two treatments (despite being slightly negative under control and positive under high night temperature), and were not significantly different from zero (*P*>0.05). Similarly, no significant trends were observed for *A*_net_^25^ when expressed per dry mass (*A*_net_DM_), fresh mass (*A*_net_FM_) or nitrogen (*A*_net_N_, except for the control) Supplementary Figure S2). Leaf *A*_net_N_ under control treatment declined (adjusted *R*^2^□=□0.51, *p* < 0.05) with Year of Release (Supplementary Figure S2c). This indicates potential decoupling of photosynthesis from leaf nitrogen content in modern cultivars under non-warm night conditions.

**Figure 1.**
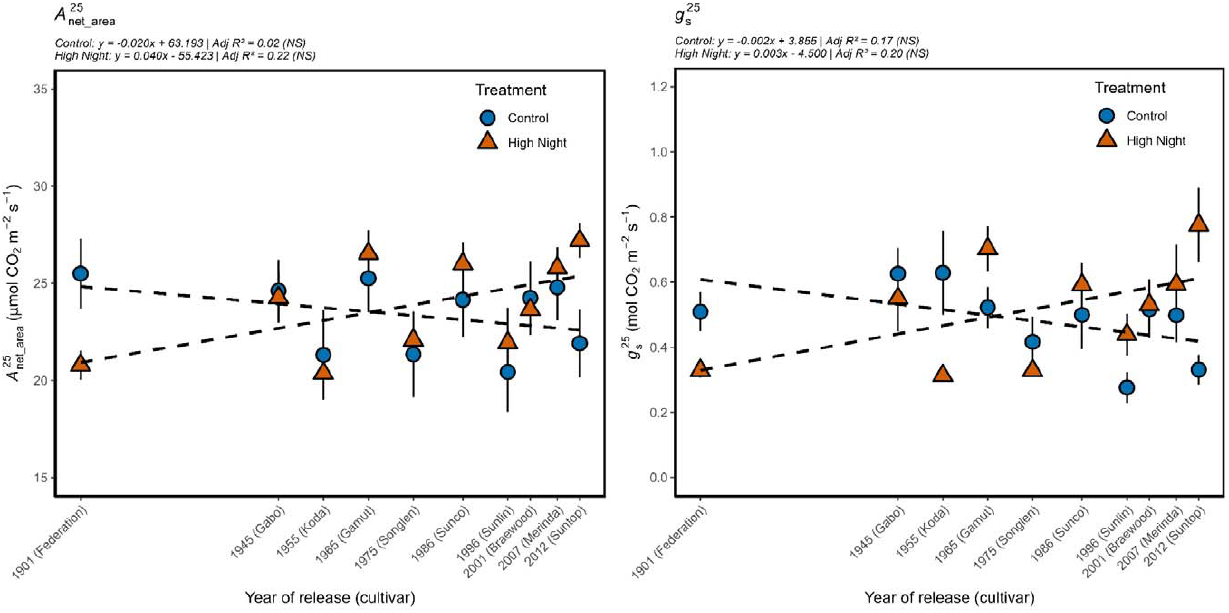
Net rates of photosynthetic CO_2_ assimilation per unit leaf area (*A*_net_area_, µmol CO_2_m^2^s^−1^; left panel) measured at 25 °C; and stomatal conductance (*g*_s_, mol CO_2 m_^−2^s^−1^; right panel), also measured at 25 °C, of Australian wheat cultivars released between 1901 and 2012. These were treated to night temperatures of 12 °C (control, blue circles) or 22 °C (high night, red triangles). Adjusted *R*^2^ are indicated at the top of each panel. Where regression slope is not significantly different from zero this is indicated by black dotted lines and NS next to the corresponding adjusted *R*^2^. Error bars are s.e.m.s *n* = 2-6.

### 3.2 Photosynthetic capacity: V_**cmax**_**and J**_**1500**_**declined in modern cultivars under warm nights but TPU has been stable across breeding history**

To assess the mechanism underpinning *A* _net_^25^ under night temperature treatments among the cultivars (or across Year of Release), *V*_cmax_, *J*_1500_ and TPU were determined from *A*_net_-CO_2_ response curves at 25 °C (*V*_cmax_^25^, *J*_1500_^25^ and TPU^25^). For each measured leaf, unreliable estimates of *V* _cmax_^25^, *J* _1500_^25^ and TPU^25^– with *A* _net_^25^ by chloroplastic CO_2_ (*A* _net_-*C* _c_) curves showing obvious errors – were omitted from further analyses of photosynthetic capacity (See Supplementary Figure S3 for examples). Under control conditions, *V* _cmax_^25^ showed no significant trend through time (adjusted *R*^2^= –0.01), even though it varied significantly among cultivars. For examples, mean *V*_cmax_^25^ of Federation (1901) at 204 µmol CO_2_m^2^s^−1^ was almost double the mean *V*_cmax_^25^ of Merinda (2007) at 108 µmol CO_2_m^2^s^−1^. By contrast, under high night temperature, a negative decline was observed from 1955 (Koda) under high night temperature (adjusted *R*^2^= 0.60; *p* < 0.05; Figure 2a), indicating reduced carboxylation capacity in more recent cultivars when exposed to warm nights.

**Figure 2.**
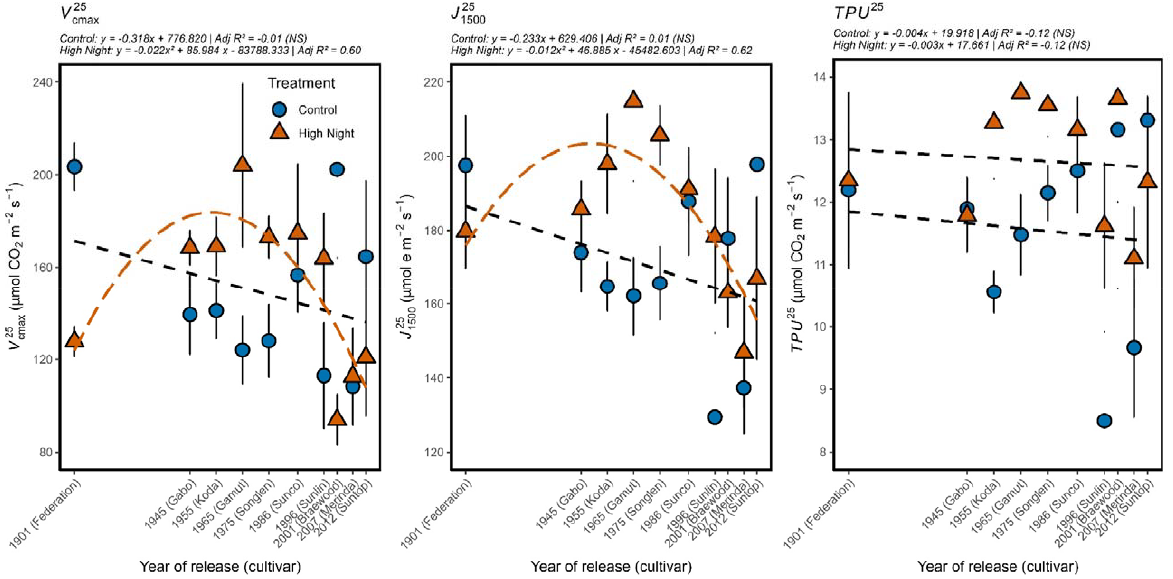
Photosynthetic efficiency traits measured at 25 °C of ten Australian cultivars released between 1901 and 2012 and treated to night temperatures of 12 °C (control, blue circles and lines) or 22 °C (high night, red triangles and lines). Maximum rate of Rubisco carboxylation (*V*_cmax_, left panel (a)), rate of electron transport (*J*_1500_, middle panel (b)), and rate of triphosphate utilisation (TPU, right panel (c)). Adjusted *R*^2^ values are indicated at the top of each panel. Where regression slope is not significantly different from zero this is indicated by black dashed lines and NS next to the corresponding adjusted *R*^2^ value. Significant slopes are presented with treatment-specific colour. Error bars are s.e.m. *n* = 2-6.

Only two cultivars significantly downregulated *V*_cmax_^25^ under warm nights, and these were Federation (1901) and Braewood (2001). Of the remaining eight cultivars, five mostly older cultivars (1945-1996) increased *V*_cmax_^25^ and three (including two new cultivars) maintained *V*_cmax_^25^ under warm nights, with the greatest increase exhibited by Gamut (1965). *J*_1500_^25^ largely followed the response patterns to *V*_cmax_^25^ under control and high night temperatures – no significant trend detected under control conditions and declining with Year of Release from 1955 (Koda) (adjusted *R*^2^= 0.62; Figure 2b). TPU^25^ exhibited little variation under either treatment (adjusted *R*^2^= –0.12 for both control and high night temperature). Mean TPU^25^ under control and high night temperatures were both 12.5 µmol CO_2_m^2^s^−1^ and their slopes were not significantly different (Figure 2c). These suggests stability in TPU^25^ capacity across breeding history. Under warm night conditions, most older cultivars had significantly higher *V*_cmax_^25^ and *J*_cmax_^25^ than newer cultivars (Figure 2a-b). The difference was largest for the cultivar Gamut (1965) for both *V*_cmax_^25^ and *J*_1500_^25^. Consequently, the *J*_1500_^25^:*V* _cmax_^25^ was highest for Gamut (1965) (Figure 3a).

**Figure 3.**
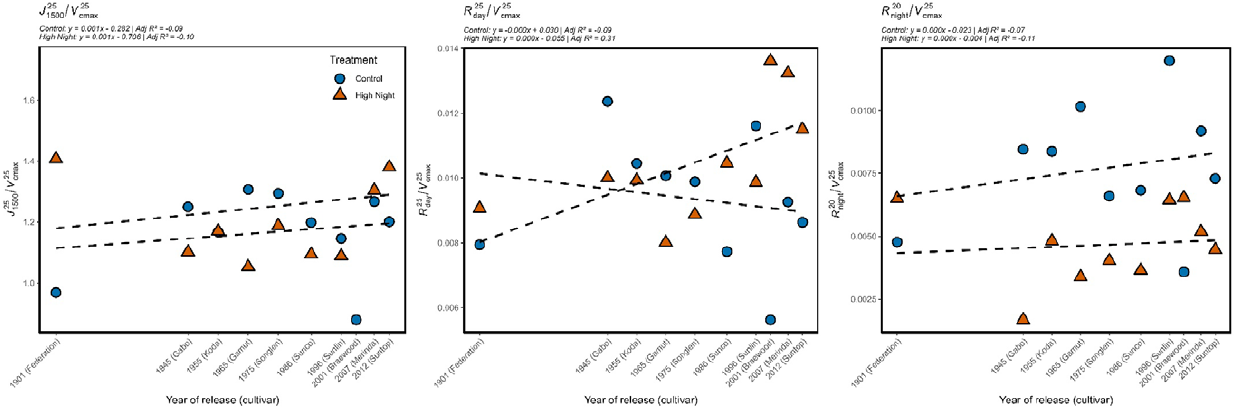
Changes in ratio of wheat leaf photosynthetic efficiency parameters (rates of electron transport, *J*_1500_ to rates of Rubisco carboxylation, *V*_cmax_) measured at 25 °C (left panel), and ratios of respiration during the day (*R*_day_) and measured at 25 °C (middle panel) and respiration at night (*R*_night_) measured at 20 °C (right panel) to *V*_cmax_ measured at 25 °C when treated to different night temperatures. Estimates were made using ten Australian cultivars released between 1901 and 2012. The wheat cultivars were treated to control night temperature (12 °C, blue circles and line) or high night temperatures (22 °C, red triangles and lines) at anthesis. *n* = 2-3.

### 3.3 Leaf respiration: warm nights enhanced day respiration but caused downregulation of night respiration

Wheat leaf *R*_day_^25^ and *R*_night_^20^ showed different trends between temperature treatments and varied among cultivars (Figure 4). On one hand, *R*_day_^25^ exhibited a significant linear decline with Year of Release under control conditions (adjusted *R*^2^= 0.47); *R*_day_^25^ also declined with Year of Release under warm nights, but only from 1945 with Gabo (adjusted *R*^2^= 0.58). The curvilinear response of *R*_day_^25^ under high night temperature, depicted in Figure 4a, was largely driven by the oldest cultivar Federation (1901), which had the lowest *R*_day_^25^ among all genotypes. Overall, temperature-normalised rates of *R*_day_ were either maintained or increased under warm nights except for Federation (1901) which had lower *R*_day_^25^ in plants treated to warm nights. By contrast, temperature-normalised *R*_night_ showed no discernible temporal trend under either control or warm night conditions, and for most cultivars was reduced under high night temperature (Figure 4b) – suggesting that *R*_night_ thermally acclimated in most cases. At the common temperature of 25 °C, *R*_night_^25^ was higher than *R*_day_^25^ under control conditions. But the reverse was the case under high night conditions with *R*_night_^25^ being lower than *R*_day_^25^ (Supplementary Figure S4, also compare Figure 4a versus 4c). Warm nights resulted in the relationship between *R*_dark_ and *V*_cmax_ and time being changed from a negative (control) to positive (warm night) slope for *R*_day_^25^ (adjusted *R*^2^= 0.31; Figure 3b). Additionally, warm nights markedly lowered *R*_night_^20^/*V* _cmax_^25^ compared to controls (Figure 3c).

**Figure 4.**
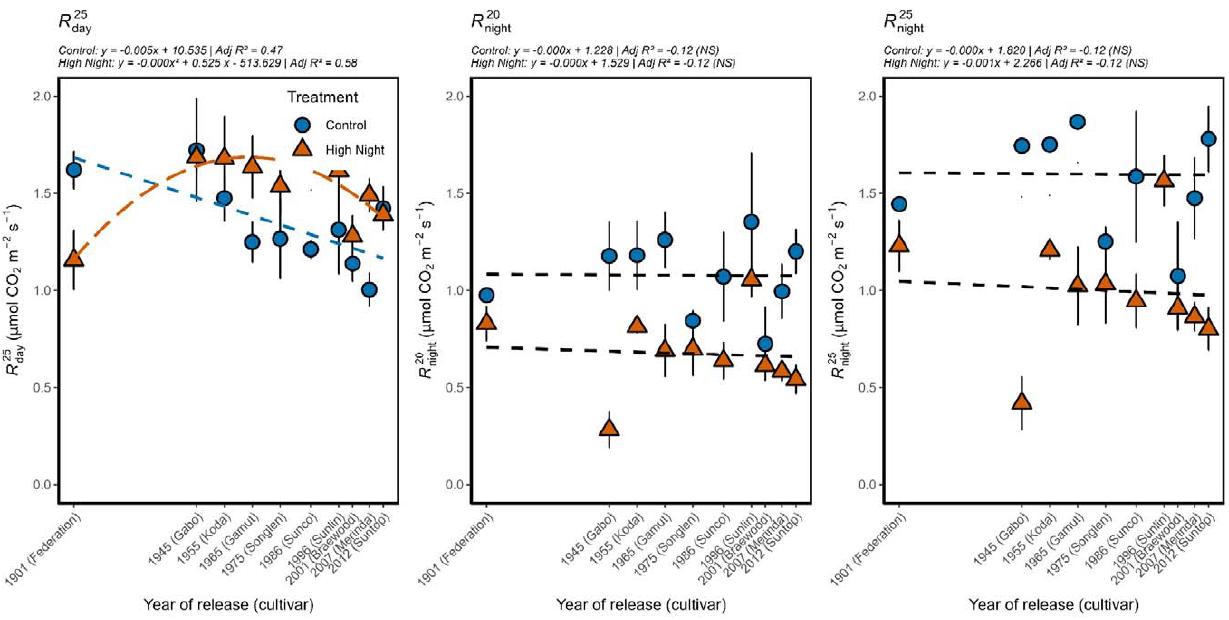
Response of wheat leaf CO_2_-based respiration of ten Australian cultivars released between 1901 and 2012 and grown at control (12 °C, blue circles and line) or high night temperatures (22 °C, red triangles and lines) at anthesis. Leaf respiration was measured during the day at 25 °C (*R*_day_^25^; left panel) and during the night at 20 °C (*R* _night_^20^; middle panel). For comparison with *R*_day_^25^ at a common measurement temperature of 25 °C, leaf respiration during the night at 25 °C (*R*_night_^25^; left panel) was estimated using the global polynomial respiration-temperature model of Heskel et al (2016). Adjusted *R*^2^ for regression lines are indicated at the top of each panel. Significant slopes are presented with treatment-specific colour and non-significant slopes by black dashed lines and NS next to the corresponding adjusted *R*^2^. Error bars are s.e.m. *n* = 2-6.

### 3.4 Leaf functional traits: Leaf nitrogen content increased but leaf mass per area remained unchanged under warm nights

Leaf N content (N_mass_ and N_area_) increased with Year of Release (i.e. was different among cultivars, *p* < 0.01), more markedly in response to warm nights (Figure 5a-b). By contrast, LMA showed no consistent temporal trend under either treatment (Figure 5a-b). Leaf N_area_ was associated either linearly or curvilinearly with photosynthetic capacity traits (*V*_cmax_^25^ and *J*_1500_^25^) under both control and high night temperature (Figure 6a-b). The relationship was strongest for *V*_cmax_^25^ (adjusted *R*^2^= 0.54) under warm nights followed by *J*_1500_^25^ (adjusted *R*^2^= 0.37), also under warm nights. These adjusted *R*^2^ represent the proportion of variations explained by the model considering the different intercepts and slopes of the night temperature treatments. At low N_area_ (~1.4 g m^−2^) and high N_area_ (>2.2 g m^−2^) *V*_cmax_^25^ and *J*_1500_^25^ were higher under warm nights than control conditions. By contrast, leaf *R*_day_ declined with increasing N_area_ both under control and warm nights, while *R*_night_ exhibited a non-linear response to N_area_ under warm night (Figure 6d-e).

**Figure 5.**
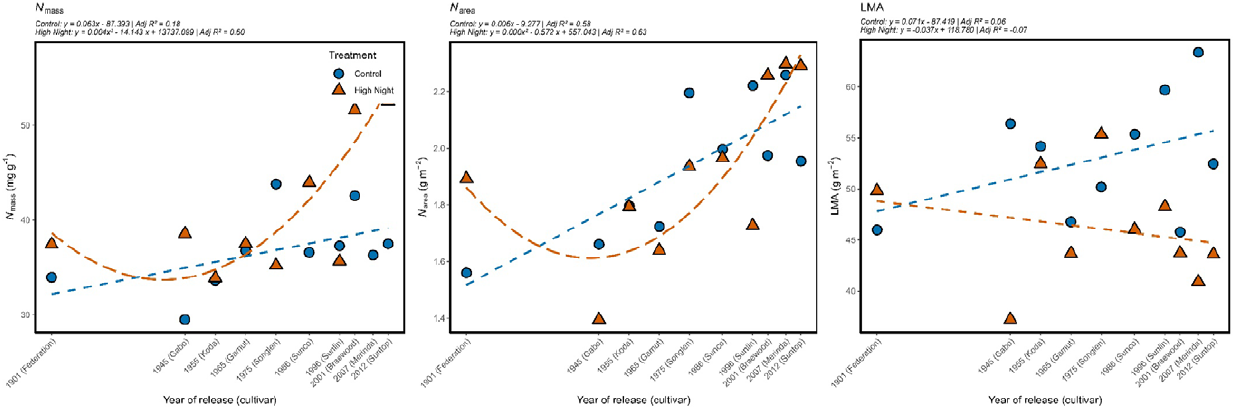
Changes in wheat leaf functional traits (nitrogen per unit leaf mass, *N*_mass,_ left panel (a); nitrogen per unit leaf area, *N*_area_, middle panel (b); and leaf mass per unit area, LMA, right panel (c)) of ten Australian cultivars released between 1901 and 2012. The wheat cultivars were treated to control night temperature (12 °C, blue circles and line) or high night temperatures (22 °C, red triangles and lines) at anthesis. *n* = 2-6.

**Figure 6.**
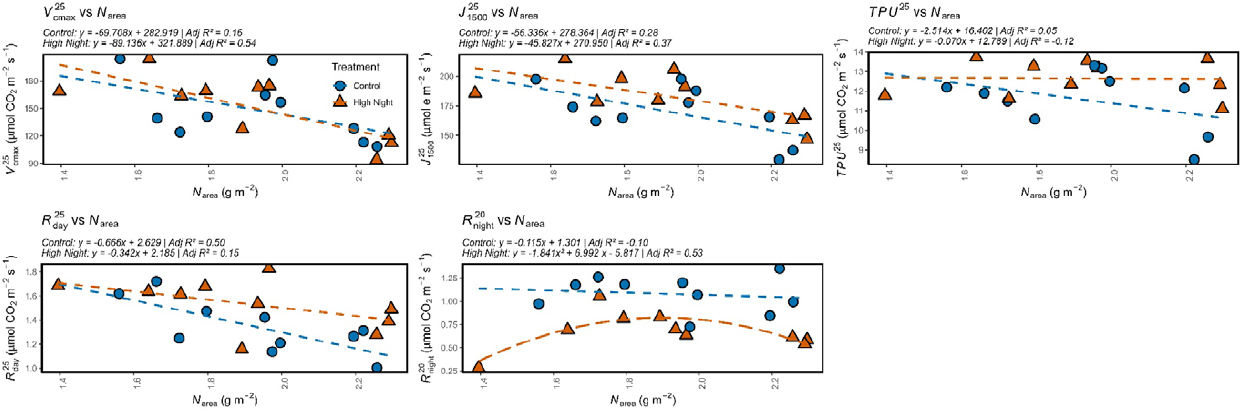
Relationships of wheat leaf photosynthetic capacity (maximum rates of Rubisco carboxylation, *V*_cmax_^25^ (a); rates of electron transport, *J*_1500_^25^ (b); rates of triphosphate utilisation, TPU^25^ (c)), respiration during the day (*R*_day_, (d)), respiration at night (*R*_night_, (e)) with nitrogen per unit leaf area (N_area_) of ten Australian cultivars released between 1901 and 2012. The wheat cultivars were treated to control night temperature (12 °C, blue circles and line) or high night temperature (22 °C, green triangles and lines) at anthesis. *n* = 2-6.

## 4.0 Discussion

Globally and in Australia wheat yield has increased over the last several decades due to advances in breeding and improved agronomic management (Fischer et al., 2014; Richards et al., 2014). It is assumed that conventional breeding and selection for yield increases has inadvertently selected for enhanced photosynthesis (*A*_net_ measured at a common temperature, e.g. at 25 °C; *A*_net_^25^) or lower temperature-normalised *R*_dark_. However, with rising night temperatures, causing a drag on wheat yield, it is unclear how well these physiological traits acclimate to warm nights. In our study, leaf *A*_net_^25^ showed no systematic change with Year of Release, except when expressed per unit N, where older cultivars exhibited greater *A* _net_^25^ under warm nights (Figure 1c). However, these trends were not mirrored in the biochemical components that underpin C_3_ photosynthetic capacity, with modern cultivars exhibiting reduced *V*_cmax_^25^ and *J*_1500_^25^ under warmer nights. This pattern contrasts with the hypothesis that conventional breeding has inadvertently selected for enhanced photosynthesis. Leaf *R*_dark_ measured during the day at 25 °C (*R*_day_^25^) and during the night at 20 °C (*R*_night_^20^) or estimated at 25 °C (*R*_night_^25^) showed consistently divergent responses to warming: leaf *R* _day_^25^ declined with Year of Release but was stimulated under warm nights, whereas leaf *R*_night_^20^ or *R*_night_^25^ remained unchanged across time but was significantly downregulated under warm nights. These contrasting thermal sensitivities suggest differential metabolic control of *R*_day_ and *R*_night_, possibly reflecting circadian regulation, substrate availability, or partitioning of energy demands during light and dark periods. Understanding the mechanisms driving these divergent responses offers a pathway to inform targeted breeding strategies for wheat cultivars better adapted to future climates.

### 4.1 Limited gains to date in photosynthesis rate but stable response to warm nights might enhance wheat yield resilience

Historical yield gains in Australia do not appear to have been underpinned by systematic improvements in leaf photosynthesis across cultivars released between 1901 to 2012 under control night temperatures. This suggests that breeding strategies focused on yield and stress resilience have not inadvertently selected for increased photosynthetic rates or capacity. Nonetheless, enhancing photosynthesis remains a viable strategy for boosting wheat productivity under future climates (Burgess et al., 2023; Croce et al., 2024; Gracia-Romero et al., 2023; Yin et al., 2022), especially by maximising the efficiency of converting intercepted radiation into biomass (Garcia et al., 2023). Several physiological strategies have been proposed to improve photosynthetic efficiency, including: (i) increasing the catalytic performance of Rubisco (ribulose-l,5-bisphosphate carboxylase/oxygenase) and enhancing electron transport and mesophyll conductance (Parry et al., 2011; Prins et al., 2016; Whitney et al., 2011; Wu et al., 2019; Yin et al., 2022); (ii) exploiting photosynthetic contributions of non-foliar tissues including spikes, tillers and awns (Simkin et al., 2020; Vicente et al., 2024); (iii) modifying canopy architecture to improve light interception and increase radiation use efficiency (Guarin et al., 2022); and (iv) engineering photorespiratory pathways to reduce CO_2_ losses (Garcia et al., 2023; Hagemann & Bauwe, 2016; South et al., 2018). Modelling estimates suggest that simultaneous improvements in Rubisco activity, electron transport and mesophyll conductance could increase wheat yield by 5.3-12.1% (Wu et al., 2019). However, others caution that gains in photosynthesis are unlikely to translate to higher yields unless coupled with coordinated improvements in nitrogen acquisition, assimilation, and remobilisation to grains (De Souza et al., 2023; Sinclair et al., 2023; Sinclair et al., 2019).

Acclimation of photosynthesis to warming typically involves enhancement of *A*_net_ at a common temperature in warmer-grown plants and increase in the optimum temperature (*T*_opt_) of *A*_net_ at the warmer growth temperature (Berry & Bjorkman, 1980; Way & Yamori, 2014). In our study, the stable *A*_net_^25^ under warming across most cultivars (Figure 1a, b) could be due to limited sensitivity to night temperature in the low 20 °C range. These findings are consistent with prior studies in wheat: Coast et al. (2024) observed *A*_net_^25^ maintenance or increases under nights of 20 and 25 °C in three out of four genotypes; and Impa et al. (2019) reported stable *A*_net_^26^ in five of six winter wheat genotypes under day/night temperature treatments of 26/23 °C. In the present study, where the night warming magnitude was 8.7 °C, nine of ten cultivars were not negatively affected by warm nights. But the nine cultivars also did not show typical acclimation of photosynthesis to higher temperature – i.e. constructive acclimation, which is described by Way & Yamori (2014) as both increase in *A*_net_^25^ and T_opt_ of *A*_net_ at the warmer growth temperature. The exception was the earliest released cultivar, Federation (1901), which consistently exhibited a decline in *A* _net_^25^ under warming i.e. detractive acclimation, regardless of whether it was expressed per unit area, mass or nitrogen. These results demonstrate that while long-term selection for yield has not led to a steady improvement in *A*_net_^25^ (or typical acclimation) it has maintained photosynthesis rates under warm nights in modern cultivars. Given that rising night temperature is a hallmark of climate change (IPCC, 2013), the ability of wheat leaves to at least maintain photosynthetic performance under warm nights may represent an important physiological mechanism contributing to yield stability in future climates.

### 4.2 Decreasing photosynthetic capacity in modern cultivars but different cultivar sensitivities to warm nights provide opportunity for increasing photosynthesis

Under saturating light and adequate water supply, photosynthesis in C□plants is limited by the kinetics of carbon fixation (*V*_cmax_) and RuBP regeneration via maximum electron transport rate (*J*_max_) (Farquhar & Sharkey, 1982; Farquhar et al., 1980). These two processes underpin photosynthetic capacity and are sensitive to thermal and genotypic variation. In our study, photosynthetic capacity showed no clear trend with cultivar Year of Release under control night temperatures, consistent with reports from historical soybean germplasm (Koester et al., 2016). However, under warm nights, photosynthetic capacity declined with increasing Year of Release midway through the century, and most cultivars exhibited higher *V*_cmax_^25^, *J* _1500_^25^ and TPU^25^ under warming. The exception being Federation, released in 1901, and Suntop, released in 2012; Figure 2) – which showed higher *V*_cmax_^25^ and *J*_1500_^25^ under control conditions (Figure 2a). These results show that there are modern cultivars able to maintain or improve photosynthetic capacity under warm nights, i.e. thermally acclimate to warm nights. Although the cool-night preference of these modern cultivars may align with a hypothesis of greater N allocation to photosynthetic machinery at lower temperatures, their absolute *V*_cmax_^25^ and *J*_1500_^25^ values were lower than those of historical cultivars. Importantly, these reductions were not associated with decreased leaf N, which increased with cultivar Year of Release and warm night treatment (Figure 5a-b). This implies a possible shift or lower proportional allocation of nitrogen to photosynthetic proteins such as Rubisco or components of the electron transport chain in modern cultivars. These results are consistent with earlier reports that warming reduces *V*_cmax_ through changes in protein allocation rather than total nitrogen (Crous et al., 2018; Dusenge et al., 2020; Scafaro et al., 2017). They also align with recent meta-analysis showing reduced *V*_cmax_^25^ in warm-grown plants (Crous et al., 2022; Wang et al., 2020). The results also support the idea that selection for yield has not coincided with increased photosynthetic capacity, and may have inadvertently favoured lines with lower biochemical capacities (Driever et al., 2014) under warming. However, it is not unlikely that modern wheat cultivars can maintain *A*_net_^25^ at higher night temperatures even with less protein invested in the photosynthetic machinery, because they have more efficient photosynthetic processes. This could be beneficial for reducing *R*_dark_ (in particular *R*_night_) which would translate to less maintenance respiration required for expensive Rubisco protein turnover and synthesis.

The differing thermal sensitivities of *V*_cmax_^25^ and *J*_1500_^25^ suggest underlying biochemical divergence. Electron transport processes are more dependent on thylakoid membrane structure and redox balance, rendering *J* _1500_^25^ more vulnerable to high temperatures (Yamori et al., 2014). Rubisco carboxylation, while also temperature-sensitive, may be comparatively more buffered due to its enzymatic activation properties. The distinct trajectories of *V*_cmax_^25^ and *J*_1500_^25^ under warm nights further manifested in increased *J*_1500_^25^/*V* ^25^ ratios in modern cultivars, suggesting a decoupling of RuBP regeneration and carboxylation under warming. By contrast, TPU^25^ remained stable across treatments and genotypes, suggesting that TPU was not limiting under warm night stress. This is consistent with other studies reporting TPU stability under moderate stress (Wang et al., 2020), and highlights early steps of the Calvin cycle as key targets for improving photosynthetic resilience to warming.

### 4.3 Wheat leaf R_**night**_, **but not R**_**day**_, **acclimates to moderate warm nights**

A widely proposed theory for yield loss due to warm nights has been stimulation of dark respiratory CO_2_ release, which may be coupled with reduced photosynthetic capacity. In response to sustained or chronic increase in temperature, plants acclimate respiration by reducing their respiration rate at a given measurement temperature (Atkin & Tjoelker 2003; Crous et al., 2011). In our study, warm nights consistently downregulated temperature-normalised rates of *R*_night_ while stimulating *R*_day_, indicating that nocturnal respiration acclimated to warming, whereas daytime respiration did not. These divergent responses are consistent with previous observations in wheat: *R*_night_ acclimation to night warming (Impa et al., 2019), and no acclimation of *R*_day_^25^ under high night temperature regimes (Posch et al., 2022). In Posch et al. (2022), the only decrease in *R*_day_ which occurred under an extreme night temperature (25□°C) and minimal diurnal range (26/25□°C), suggesting that treatment severity may drive atypical responses, was however not significant (i.e. no acclimation). In the present study, the divergent responses of temperature-normalised *R*_day_ and *R*_night_ to nocturnal warming resulted in an overall reduction in respiratory CO□release (~12%), as the stimulation of *R*_day_ was insufficient to offset the downregulation of *R*_night_. *R*_day_^25^ would have required a 31% increase to compensate fully.

Our study agrees with earlier reports of acclimation of *R*_night_ to night warming observed in other plants/crops treated to moderate night temperatures (low-to mid-20 °C) including rice (Peraudeau et al., 2015), soyabean (Djanaguiraman et al., 2013), loblolly pine - (*Pinus taeda* L.; Will, 2000) and *Stipa krylovii* Roshev (Chi et al. 2013). Chi et al. (2013) also reported that leaves of *Stipa krylovii* did not acclimate *R*_day_ to nocturnal warming. However, our results contradict results of increased *R*_night_ to warm nights (i.e. limited to no acclimation of temperature-normalised rates of *R*_night_) reported by others for cotton (Loka & Oosterhuis, 2010), maize (Kettler et al., 2022), sorghum and sunflower (Manunta & Kirkham, 1996), rice (Bahuguna et al., 2016), and soyabean (Djanaguiraman et al., 2013). The differences in acclimation responses might be due to imposition of very high night temperatures (>25 °C) or little diurnal amplitude (e.g. 26/25 °C, day/night regime).

The uncoupling of *R*_night_ and *R*_day_ from night temperature suggests that *R*_night_ acclimation may be regulated independently of *R*_day_. This likely reflects diurnal variation in leaf metabolic status, demand for respiratory products, and source of substrate supply used in the mitochondria for respiration. At night, sucrose is derived from circadian-controlled synthesis of remobilized transitory starch degraded in the chloroplast (Smith & Stitt, 2007; Smith & Zeeman, 2020), whereas in the day sucrose is synthesized from triose phosphates produced by the Calvin-Benson-Bassham cycle. These contrasts imply that *R*_day_, constrained by its coupling with photosynthesis, may be less free to downregulate, under nocturnal warming.

Respiratory acclimation has been linked to foliar nitrogen, reflecting the leaf N-respiratory scaling relationship and dependence of respiration on nitrogen-rich enzymes and cofactors which make up a large proportion of total N in leaves (Abadie et al., 2017; Atkin et al., 2015; Atkin & Tjoelker, 2003; Tjoelker et al., 1999). In our study, leaf N scaled positively with *R*_night_ but weakly and negatively with *R*_day_ under warm nights, further highlighting their divergent responses. The coupling of leaf N and *R*_night_ suggests enzyme limitation of respiratory capacity (Ryan et al., 1996) and highlights the potential use of leaf N as a predictor of *R*_night_ in crop and Earth system models (Atkin et al., 2015; Fisher et al., 2014). However, leaf N accounted for only about half of the variation in *R*_night_ responses, implying that other factors also contribute. Factors such as substrate availability (Covey-Crump et al., 2002), changes in soluble carbohydrate concentrations (Atkin et al., 2000b), altered enzyme capacity (Atkin & Tjoelker, 2003), and differences in the temperature sensitivity of enzymatic steps regulating day versus night respiration.

Alternatively, given that *V*_*c*max_ controls the supply of respiratory substrates (O’Leary et al., 2019) and that respiratory ATP is required to maintain protein turnover in the photosynthetic apparatus (Dusenge et al., 2021), acclimation of *R*_night_ may reflect a strategy to sustain optimal photosynthetic capacity (Sturchio et al., 2021; Wang et al., 2020). This interpretation is consistent with our finding that decreased *R*_night_ coupled with reduced but more efficient *V*_cmax_ and *J*_max_, especially in modern cultivars, resulted in maintained *A*_net_^25^. However, these patterns are not consistent with typical photosynthetic acclimation (Way & Sage, 2008; Way & Yamori, 2014), emphasising the need for further work to disentangle the regulatory links between *R*_day_ and photosynthetic capacity. Whether *R*_night_ acclimation in wheat represents Type I or Type II acclimation (Atkin & Tjoelker, 2003) was not examined in our study and remains to be tested, but our findings indicate that *R*_night_ exhibits greater plasticity and thermal sensitivity than previously recognised.

By contrast, the lack of *R*_day_ acclimation is probably linked to its coordination with photosynthetic capacity. Under warm nights, the ratio *R*_day_:*V*_cmax_ increased with year of release, suggesting that higher daytime respiratory demand may have depleted soluble carbohydrate pools, thereby constraining catalytic activity and limiting thermal downregulation of *R*_day_.

### 4.4 Towards improved parameterisation of respiration in crop and Earth system models

Our findings, that *R*_night_ acclimates to moderate warm nights whereas *R*_day_ do not, have important implications for crop and Earth system models and the reliable estimation of terrestrial carbon emissions. Most models assume that *R*_day_ (after 30 min of dark exposure during the day to avoid the post-illumination photorespiratory CO_2_ burst and light-enhanced dark respiration) equates to *R*_night_ (Atkin et al., 2000a; Butler et al., 2021; Huntingford et al., 2017; Padmasree et al., 2002) – conflating *R*_day_ and *R*_night_ responses. While *R*_day_ can be similar to *R*_night_ in some situations (Florez-Sarasa et al., 2012), there is growing evidence that the two differ (Gessler et al., 2009, 2017; Fan et al 2024; Zheng et al., 2025). As such, even though both are assessed under dark conditions, their thermal sensitivities are probably different (Bruhn et al., 2022; 2024b; Zheng et al., 2024). Our results therefore highlights that models reliant on *R*_day_ measured in darkness as a proxy for *R*_night_ measured in darkness would lead to overestimation of night-warming-induced respiratory carbon losses over long periods. This emphasizes the need for a more complete understanding of the degree to which *R*_night_ and *R*_day_ responses to warm nights differ within and among plant species. Improving parameterisation of Earth system models will require dynamic temperature response functions that explicitly distinguish between *R*_day_ and *R*_night_ and incorporate their capacity for acclimation.

### 4.5 Future research suggestions

Research is required to identify quantitative trait loci (QTLs) associated with *R*_night_, *V*_cmax_ and *J*_max_ under warm night conditions. This will be critical for uncovering the genetic basis of acclimation and enabling marker-assisted selection. Progress in breeding wheat cultivars with enhanced tolerance to warm nights can be supported by understanding the genetic architecture underlying respiratory and photosynthetic traits, particularly *R*_night_. While respiration is a complex/polygenic trait influenced by many loci with small effects and strongly modulated by environmental conditions, recent studies suggest that it is nonetheless genetically tractable. For example, wheat leaf respiration has been shown to be under both genetic and environmental control (Gaju et al., 2025), and genome-wide association studies (GWAS) are beginning to reveal the underlying complexity of respiration in wheat and other crops (Guo et al., 2021; Bulut et al. 2023).

Research is also required to characterise the type and drivers of *R*_night_ acclimation to warm nights. Attention should also be given to improving the quantification and representation of *R*_night_ temperature responses in crop and Earth system models (Bruhn et al., 2022; 2024a; 2025), and to ascertaining whether the thermal sensitivities of *R*_night_CO_2_-efflux and O_2_-uptake are coordinated or decoupled (Coast et al., 2021; Bruhn et al., 2024b). Simultaneous measurements of both *R*_night_ respiratory fluxes would allow estimation of respiratory quotient (RQ - the ratio of moles of CO_2_ released to moles of O_2_ consumed) and indicate the substrate being oxidized for mitochondrial respiration. Changes in RQ can reveal substrate use patterns and shifts in respiratory metabolism under high night temperature stress. Future research could also focus on understanding whether the patterns seen for *V*_cmax_^25^ (i.e. decreasing under warm nights in modern cultivars but probably more efficient) is held more widely. This may be related to the abundance of and activation state of Rubisco. Also to be explored is why *A*_net_ is insensitive to moderate night temperatures in contrast to *V*_cmax_ and *J*_max_.

These research directions are relevant not only to wheat but also to other economically important cereal crops that are highly sensitive to night warming, including rice, barley and sorghum (Peng et al., 2004; García et al, 2015, 2016; Prasad & Djanaguiraman, 2011). The trait-based insights from this work provide a physiological foundation for breeding crops with improved resilience to nocturnal warming. By targeting leaf gas exchange traits, in particularly *R*_night,_ future breeding efforts may enhance energy use efficiency and yield stability under climate change.

## 5.0 Conclusion

Wheat breeding over the past century in Australia, which has focused on selection for higher yields, has inadvertently maintained *A*_net_^25^ despite declines in *V*_cmax_ and *J*_1500_ under warm night conditions. Warm nights induced divergent responses in leaf respiration: while temperature normalised *R*_day_ was stimulated, *R*_night_ acclimated (i.e. temperature normalised rates were lower under warm nights), exhibiting a downregulation of respiratory CO_2_ release. The reduction in *R*_night_ is likely driven by reduced demand for maintenance respiration, potentially reflecting lower investment in Rubisco protein turnover and synthesis. The capacity for wheat *R*_night_ acclimation to warm nights has been largely overlooked, despite its potential importance in reducing respiratory carbon loss to the atmosphere and mitigating yield declines under climate change.

## Acknowledgements

We acknowledge and celebrate the First Australians on whose traditional land this research was undertaken, and pay our respect to their elders’ past, and present. P.R.S was supported by a UNE International Postgraduate Research Award, Future Food Systems CRC Top-Up scholarship, and a Sally Muir Agricultural Research Award. We thank Dr Craig Johnson, Calista McLachlan, and Benjamin Jones for technical assistance.

## Author contributions

P.R.S, A.P.S. and O.C. planned and designed the experiment. R.J.T. and R.M.T provided the seeds. P.R.S. performed the experiment, analysed the initial data and wrote the first version of the manuscript. All authors contributed to reviewing and editing subsequent versions of the manuscript.

## Data and code availability

The data and some R codes used for this study are available as supplementary information.

---

**Supplementary Table 1.**
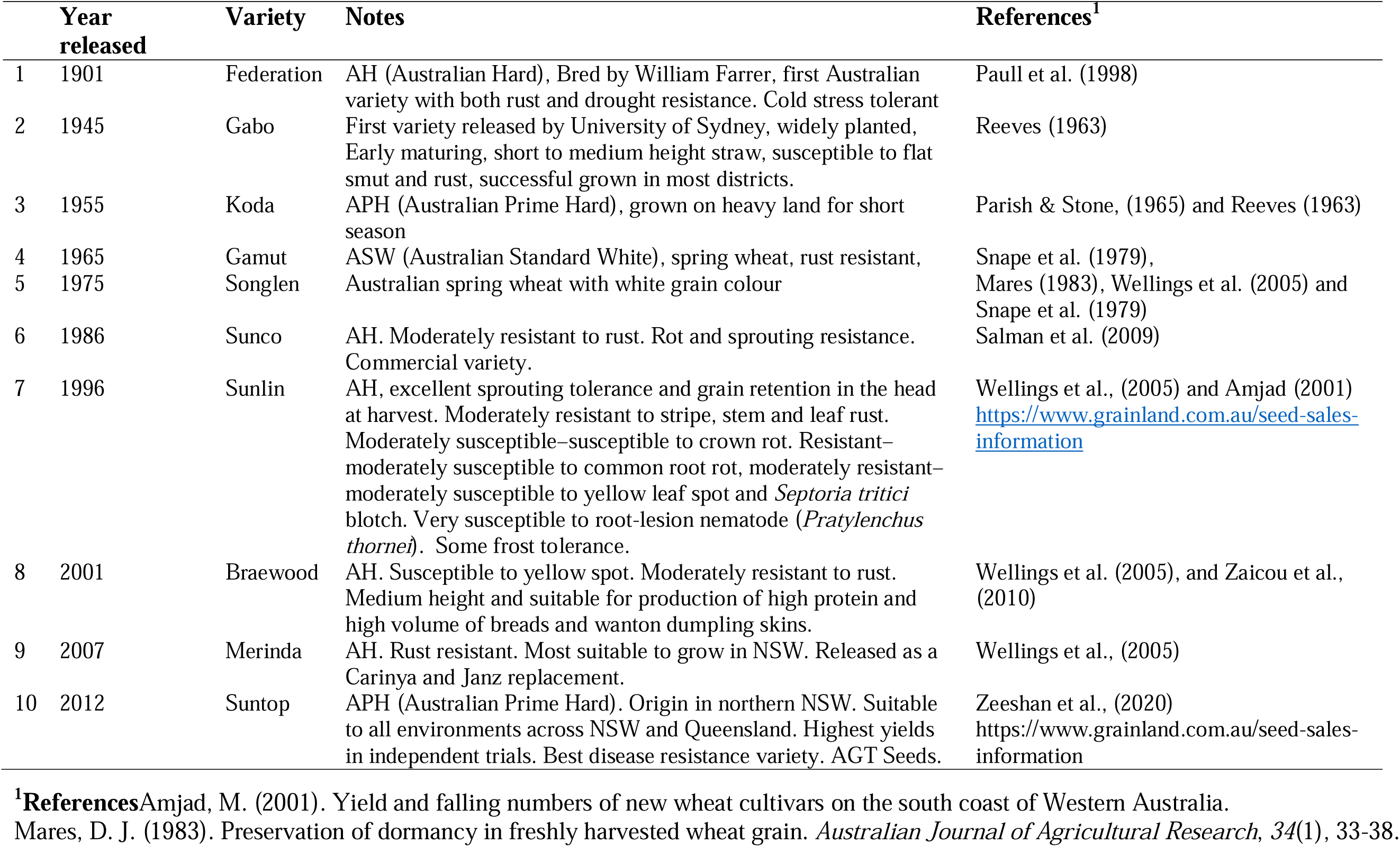
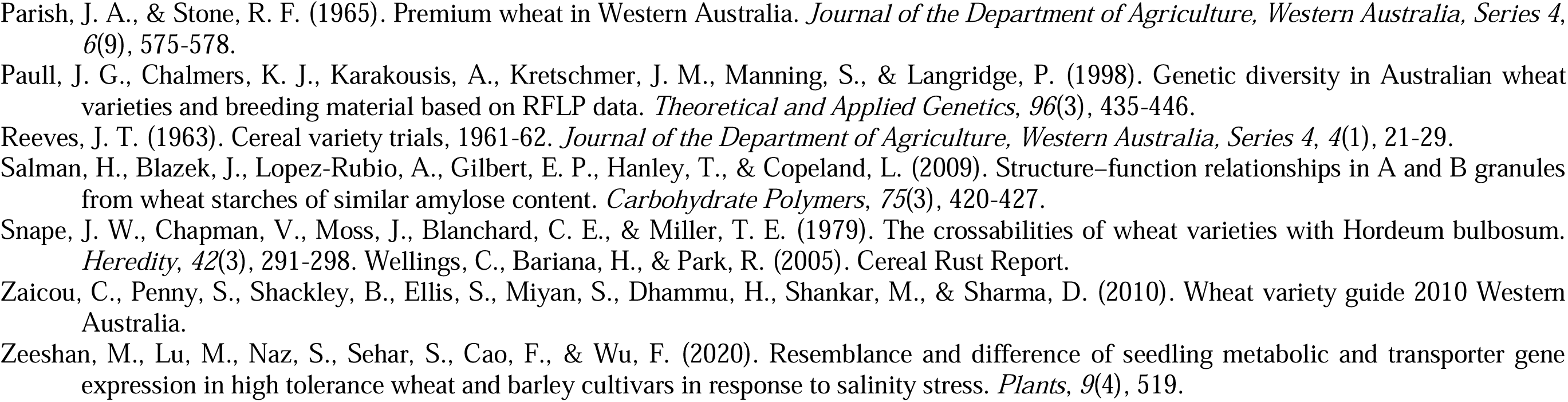
Pedigrees, year of introduction, and susceptibility traits of the wheat cultivars used in this study.

**Supplementary Figure S1.**
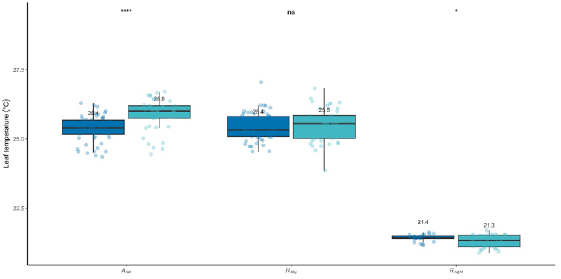
Leaf temperatures during measurements of net CO_2_ assimilation at 25 °C (*A*_net_), daytime leaf respiration at 25 °C (*R*_day_) and night leaf respiration at 20 °C (*R*_night_) for control (blue boxplots) and high night temperature (green boxplots) plants. Central lines represent the medians; numbers indicate means across cultivars; points show cultivar-level observations. Boxes indicate interquartile ranges (IQR), whiskers extend to 1.5× IQR. Differences were assessed using Welch’s two-sample t-tests. Significance annotations of paired temperature treatment comparisons based on Welch’s t-test are indicated directly over boxplots as ***, * and ns for *P*<0.001, *P*<0.05 and not significant (*P*>0.05), respectively. *n* = 2-6.

**Supplementary Figure S2.**
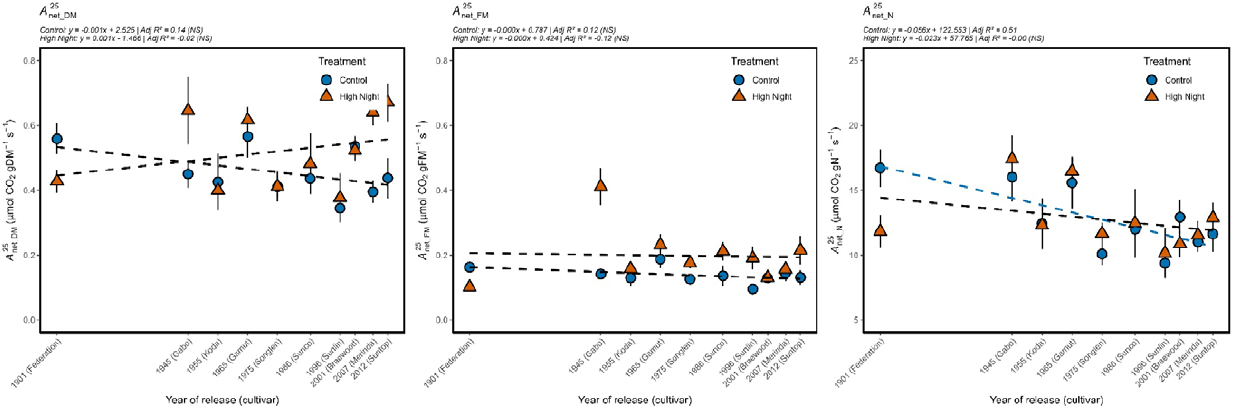
Net rates of photosynthetic CO_2_ assimilation per unit leaf dry mass (*A*_net_DM_; left panel); fresh mass (*A*_net_FM_; middle panel); and nitrogen content (*A*_net_N_; right panel) measured at 25 °C of Australian wheat cultivars released between 1901 and 2012. Cultivars were treated to night temperatures of 12 °C (control, blue circles) or 22 °C (high night, red triangles). Adjusted *R*^2^ are indicated at the top of each panel. Where regression slope is not significantly different from zero this is indicated by black dotted lines with NS next to the corresponding Adjusted *R*^2^. Significant slopes are presented with treatment-specific colour. All measurements were done at a standard reference temperature of 25°C. Error bars are s.e.m. *n* = 2-6.

**Supplementary Figure S3.**
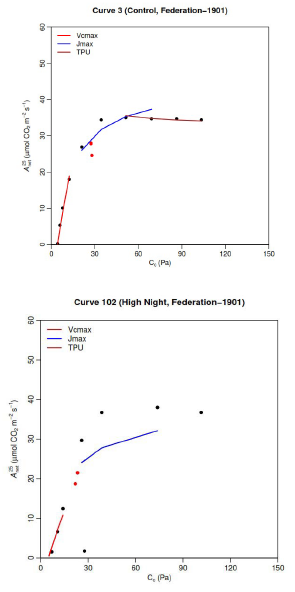
Examples of good (top panel) and bad (bottom panel) plots of net rates of photosynthetic CO_2_ assimilation measured at 25 °C (A_net,_µmol CO_2_m^−2^s^−1^;) at different intercellular CO_2_ concentration (*C*_i_, Pa). *C*_i_ was converted to chloroplastic CO_2_ concentration (*C*_c_, Pa) based on von Caemmerer and Evans (2015). Black and red dots are actual observations. Data from good plots were used for determination of photosynthetic parameters (*V*_cmax_, red lines; *J*_1500_, blue lines; and TPU, brown lines) at 25°C whereas those from bad plots were omitted from all subsequent analyses in this our study.

**Supplementary Figure S4.**
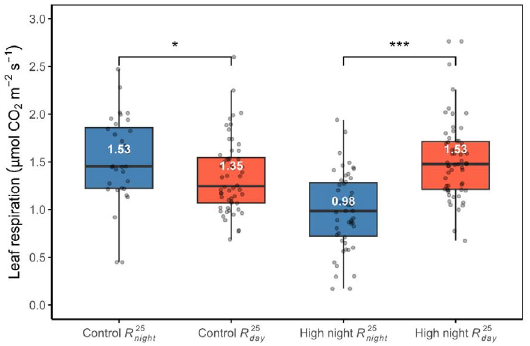
Rates of leaf respiration at 25□°C either measured during the day (*R*_day,_ red) at 25 °C or estimated from rates measured at 20□°C during the night (*R*_night_, blue) using the global polynomial respiration-temperature model of Heskel et al. (2016). Within each treatment, *R*_night_ was significantly higher than *R*_day_ under control conditions (*p*□<□0.05), whereas the reverse pattern was observed under high night conditions (*p*□<-0.001). Central lines represent the medians; bold white numbers indicate treatment means; points show cultivar-level observations. Boxes indicate interquartile ranges (IQR), whiskers extend to 1.5× IQR. Differences were assessed using Welch’s two-sample t-tests. *N* = 2-6.

